# Pan-tissue and -cancer analysis of ROR1 and ROR2 transcript variants identify novel functional significance for an alternative splice variant of ROR1

**DOI:** 10.1101/2021.11.24.469652

**Authors:** M. John, C. Ford

## Abstract

**Background:** ROR1/2 are putative druggable targets increasing in significance in translational oncology. Expression of ROR1/2 mRNA and transcript variants has not been systematically examined thus far.

**Methods:** ROR1/2 transcript variant sequences, signal peptides for cell surface localisation, and mRNA and transcript variant expression were examined in 34 transcriptomic datasets including 33 cancer types and 54 non-diseased human tissues.

**Results:** ROR1/2 have four and eight transcript variants respectively. ROR1/2 mRNA and transcript variant expression was detected in various non-diseased tissues. Our analysis identifies predominant expression of ROR1 transcript variant ENST00000545203, which lacks a signal peptide for cell surface localisation, rather than the predicted principal variant ENST00000371079. ENST00000375708 is the predominantly expressed transcript variant of ROR2.

**Conclusion:** ROR1/2 expression in healthy human tissues should be carefully considered for safety assessment of targeted therapy. Studies exploring the function and significance of the predominantly expressed ROR1 transcript variant ENST00000545203 are warranted.

## Introduction

Receptor tyrosine kinase-like orphan receptor 1 (ROR1) and ROR2 are members of the receptor tyrosine kinase family and are reported to be cell surface receptors comprising of extracellular, transmembrane and intracellular domains [1]. Multiple studies have identified Wnt5A/B and Wnt16 as cognate ligands for the receptors; hence they can no longer be considered orphans. Binding of Wnt5A (considered the primary ligand) to ROR1/2 activates non-canonical (β-catenin independent) Wnt signalling [2].

Since its identification in 1992, a pro-tumourigenic role for ROR1 signalling has been established in a growing list of both haematological malignancies (including chronic lymphocytic leukemia (CLL), mantle cell lymphoma (MCL) and (1;19) acute lymphoblastic leukemia (ALL)), and solid tumours (including ovarian, lung and chemotherapy resistant breast cancer [3–8]. A number of therapeutic strategies targeting ROR1 have reached Phase I/II clinical trials, which include a monoclonal antibody (Cirmtuzumab), antibody drug conjugates (ADC; NBE-002 and VLS-101), chimeric antigen receptor (CAR) T cell therapy and bispecific antibody (BiTE) to ROR1 and CD3 (NVG-111), for an array of malignancies [9]. Preclinical studies are also developing and evaluating additional therapeutic strategies targeting ROR1 in a growing list of cancers [9]. Since therapies inhibiting ROR1 have been shown to be well tolerated in Phase 1 clinical trials, ROR1 is now considered an elite member of the druggable genome (the subset of genes that can be pharmacologically regulated).

The current understanding of the tumourigenic role and targeted therapeutic strategies for ROR1 are based on the premise that it is a cell surface receptor activated by Wnt5A [5, 8, 10, 11]. This is based on bioinformatic prediction of the principal variant of ROR1 as a cell surface receptor as well as studies that have demonstrated cell surface expression of ROR1. However, numerous studies have consistently demonstrated cytoplasmic rather than cell surface expression of ROR1 by immunohistochemistry on both frozen and formalin-fixed, paraffin-embedded tumour samples [7, 10, 12–15]. This was confirmed by Balakrishnan et al., who developed a monoclonal antibody with careful consideration to specifically address ambiguities in previous studies [16]. The cytoplasmic localisation of ROR1 has not been adequately examined or understood to date.

Contradictory roles for ROR2 have been reported in various cancers. ROR2 mRNA expression was reported to be downregulated in primary colorectal tumours compared to normal colon epithelium in a small patient cohort and functional and *in vivo* studies pointed to a tumour suppressor role [17–19]. Our lab performed analysis of matched normal tissue and adenomas, along with analysis of colorectal tumours and cell lines and demonstrated epigenetic regulation of ROR2 expression [18]. *In vitro* experiments confirmed the epigenetic downregulation of ROR2 in colorectal cancer [19]. Conversely, ROR2 mRNA and protein expression was reported as upregulated in colorectal tumours compared to normal tissue and higher expression was associated with more severe disease and poor outcomes [20]. Similarly, contradictory studies of ROR2 in ovarian cancer have been reported [6, 21]. These inconsistencies in localisation and/or function of ROR1 and ROR2 may partly be due to differences in specificities of reagents employed for examining gene expression and function, cohort characteristics or sample processing [18]. We have previously highlighted concerns about the specificity of some commercial ROR2 antibodies [18], but this analysis has not been updated recently or expanded to include ROR1. Additionally, discrepancies may arise from the transcript variant examined in each study.

The enormous complexity of the human genome has been highlighted in recent years with the identification of multiple transcript variants for most multiexon genes (~95%) [22]. Correctly designating the functional isoform/s of a specific gene is of enormous importance because localisation and/or structure and/or function can vary dramatically between isoforms of the same gene [23]. Currently, most databases and high-throughput methods select a single transcript variant as the reference variant for each gene; however, automating this process is technically challenging with room for misidentification of functionally relevant isoforms. In some cases, the longest variant is arbitrarily selected as the reference variant. Automated annotation pipelines that have been used extensively include APPRIS (developed by the GENCODE consortium) and Matched Annotation between NCBI and EBI (MANE) [24]. APPRIS selects the transcript variant with the highest degree of conserved structural and functional motifs as the reference variant, termed the principal variant (P1) [24]. MANE Select transcript variants are those that have been annotated independently by both Ensembl and NCBI as the most biologically relevant, with matching sequences, start and end sites, 5’ and 3’ UTRs, and splice junctions. Transcript Support Level (TSL) is a methodology to describe the confidence in a given transcript model for a gene. There are five TSL categories (TSL1-5), with TSL1 having the highest and TSL5 having the lowest level of support.

Sub-cellular localisation of proteins is determined by short amino acid sequences called signal peptides (SPs), which are usually present at the amino terminus of proteins. Since these sequences are highly conserved, bioinformatic tools can identify SPs within protein sequences and predict their sub-cellular localisation [25]. ROR1 and ROR2 have been widely accepted as cell surface receptors because the isoform annotated as the reference variant for both genes have SPs that determine cell surface localisation.

Large scale RNAseq studies of non-diseased human tissues carried out by the Genotype-Tissue Expression (GTEx) project and tumour tissues by The Cancer Genome Atlas (TCGA) enable analysis of transcript variant expression in healthy tissues and human cancers [26, 27]. Since the ROR1/2 pathway is potentially both druggable and therapeutically efficacious, it is imperative to gain a clear understanding of the role this pathway plays in various cancers. Identifying the functional transcript variant/s is essential for interpreting data and for examining the role this pathway plays in disease pathophysiology.

This study explores the sequence, sub-cellular localisation and expression of various transcript variants of ROR1 and ROR2 in healthy human tissues and tumour samples. Numerous studies have reported cytoplasmic ROR1 expression and contradictory findings for the role of ROR2 in cancer; hence we sought to determine the functionally significant transcript variant that ought to be examined. We report novel functional significance for a transcript variant of ROR1 that lacks a SP for cell surface localisation, raising important questions about the mechanism of action of a gene widely considered druggable because of its cell surface localisation in cancer tissues.

## Results

### Sequence analysis of transcript variants of human ROR1 and ROR2 genes

Ensemble (release 104) annotation of the GRCh38 human reference genome assembly identified four transcript variants of ROR1 (ENST00000371079, ENST00000371080, ENST00000545203 and ENST00000482426, hereafter referred to as ROR1-v1 to ROR1-v4 respectively; Table 1). Three of the four transcript variants of ROR1 coded for proteins (ROR1-v1/2/3), all of which had complete sequences for their open reading frames (ORF), as annotated by Ensemble and the GENCODE consortium (GENCODE basic designation; Table 1). ROR1-v1/2 was categorised as TSL1 while ROR1-v3/4 was categorised as TSL5 (Table 1). Both APPRIS and MANE annotation pipelines predicted ROR1-v1 as the functional isoform of ROR1 and all of the five bioinformatic tools we employed for signal peptide prediction found a SP comprising the first 29 aa residues of this variant (Table 2). Interestingly, a SP for cell surface localisation was present in ROR1-v2 but absent in ROR1-v3 (Table 2 and Fig. 1).

**Table 1:** Transcript variants of human ROR1 and ROR2 genes. Transcript variants annotated in the GRCh38 dataset from the Genome Reference Consortium was assessed on the Ensembl genome browser (release 104). TSL: transcript support level; TSL1: transcript variants supported by a minimum of one non-suspect mRNA for every splice junction; TSL2: transcript variants supported by multiple ESTs or where the strongest supporting mRNA is flagged as suspect; TSL3: transcript variants supported by a single EST; TSL4: best supporting EST for the transcript variant is flagged as suspect; TSL5: transcript variants for which the structural model is not supported by a single transcript

| Transcript Name | Transcript ID | Transcript length (bp) | Protein length (aa) | Transcript classification | Annotation |  |  |  |  |
| --- | --- | --- | --- | --- | --- | --- | --- | --- | --- |
|  |  |  |  |  | GENCODE | Ensemble | APPRIS | MANE Select | Transcript Support Level |
| **ROR1-v1** | ENST00000371079.6 | 5858 | 937 | Protein coding | basic | Canonical | P1 | MANE Select v0.93 | TSL 1 |
| **ROR1-v2** | ENST00000371080.5 | 2305 | 393 | Protein coding | basic |  |  |  | TSL 1 |
| **ROR1-v3** | ENST00000545203.2 | 5362 | 882 | Protein coding | basic |  |  |  | TSL 5 |
| **ROR1-v4** | ENST00000482426.1 | 900 | No protein | Processed transcript |  |  |  |  | TSL 5 |
| **ROR2-v1** | ENST00000375708.4 | 4158 | 943 | Protein coding | basic | Canonical | P1 | MANE Select v0.93 | TSL 1 |
| **ROR2-v2** | ENST00000375715.5 | 2993 | 704 | Protein coding | basic |  |  |  | TSL 1 |
| **ROR2-v3** | ENST00000550066.5 | 4360 | No protein | Processed transcript |  |  |  |  | TSL 2 |
| **ROR2-v4** | ENST00000495386.5 | 664 | No protein | Processed transcript |  |  |  |  | TSL 3 |
| **ROR2-v5** | ENST00000546883.1 | 501 | No protein | Processed transcript |  |  |  |  | TSL 3 |
| **ROR2-v6** | ENST00000493846.1 | 370 | No protein | Processed transcript |  |  |  |  | TSL 3 |
| **ROR2-v7** | ENST00000476440.1 | 316 | No protein | Processed transcript |  |  |  |  | TSL 5 |
| **ROR2-v8** | ENST00000548585.2 | 334 | No protein | Retained intron |  |  |  |  | TSL 5 |

**Table 2:**
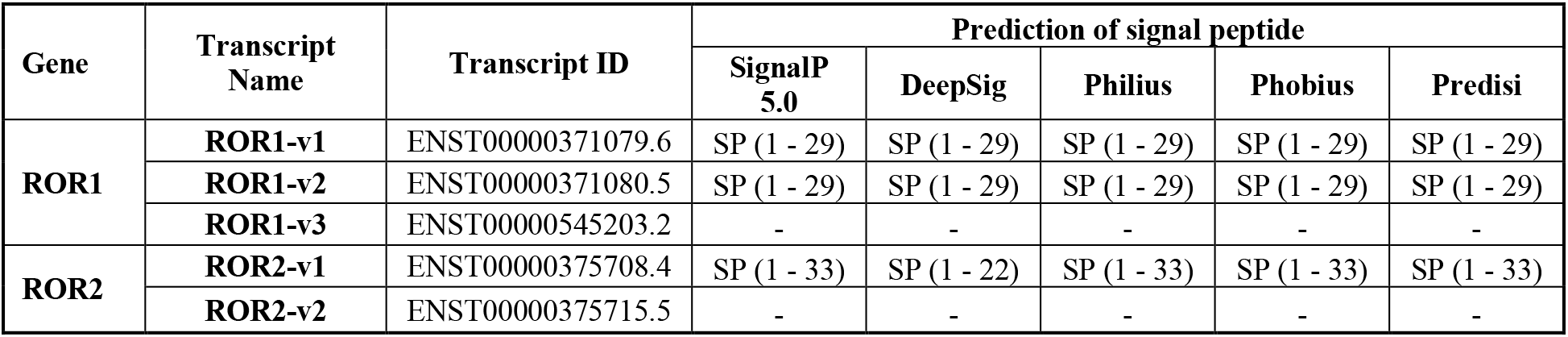
Signal peptide prediction of ROR1 and ROR2 protein coding transcript variants. Amino acid residues predicted by the SP prediction software SignalP 5.0, DeepSig, Philius, Phobius and Predisi as SPs are indicated within the parentheses.

**Figure 1:**
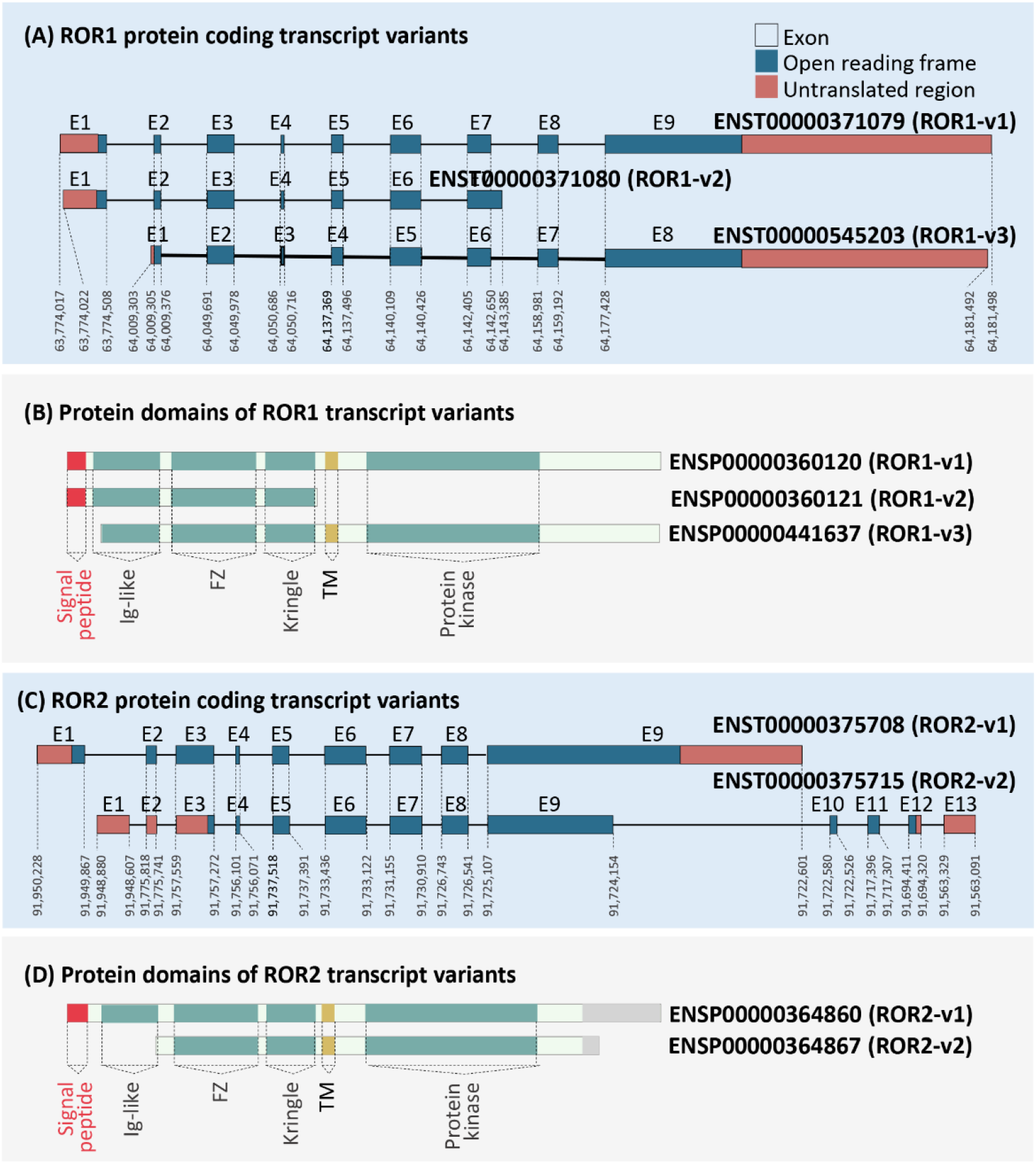
Analysis of protein coding ROR1 and ROR2 isoforms. Coding transcript and protein sequences for ROR1 (A and B) and ROR2 (C and D). Exon (E) numbers are represented above and genomic locations for ROR1 and ROR2 on chromosome 1 and 9 respectively are provided below each exon. Protein domain structures identified by uniport and bioinformatic SP prediction have been highlighted (translation IDs for the respective transcript variants are provided beside the structure).

The cDNA and protein sequences of ROR1-v1-3 were compared to identify structural and functional differences. As shown in Fig. 1A, ROR1-v1, ROR1-v2 and ROR1-v3 was comprised of nine, seven and eight exons respectively. Multiple sequence alignment established that ROR1-v3 lacked the first exon present in ROR1-v1 (supplementary data S1). Exon one and nine of ROR1-v1 included a 5’- and 3’-UTR respectively, while exon one and eight of ROR1-v3 included a 5’- and 3’-UTR respectively (Fig. 1A; supplementary data S1). The start codon for ROR1-v1 was ATG; however, a TCT codon in the second exon was predicted to be a non-canonical start codon for ROR1-v3 (supplementary data S1) and remains to be experimentally confirmed. The first two ATG codons that precede this codon were not in frame with the protein; however, a third ATG codon present after the predicted start codon was in frame with the protein sequence (supplementary data S1). Protein translation starting from the TCT codon or the first in-frame ATG codon resulted in the absence of the SP that comprises amino acid residues 1-29 of ROR1-v1 (Fig. 1B). Hence, while the protein product of transcript variant ROR1-v1 was predicted to localise to the plasma membrane, the protein product of ROR1-v3 may be retained within the cell. This possible difference in localisation of the two variants could indicate a dramatic difference in functionality.

The recently updated (May 2021) release 104 of Ensemble identified eight transcript variants for ROR2 (ENST00000375708, ENST00000375715, ENST00000550066, ENST00000495386, ENST00000546883, ENST00000493846, ENST00000476440 and ENST00000548585, hereafter referred to as ROR2-v1 to ROR2-v8 respectively; Table 1). Only two of the eight transcript variants of ROR2 coded for proteins (ROR2-v1 and ROR2-v2), both of which had complete ORF sequences, as annotated by Ensemble and the GENCODE consortium and both had TSL1 designations (Table 1 and Fig. 1C). ROR2-v1 was annotated as the MANE Select transcript variant and the principal variant (P1) by APPRIS (Table 1). Importantly, previous Ensemble releases had reported both ROR2-v1 and ROR2-v2 as equally important based on APPRIS annotation. ROR2-v1 was categorised as APPRIS P2, the designation given to two or more transcript variants when APPRIS fails to identify a single principal variant. ROR2-v2 was categorised as APPRIS ALT2, the designation given to transcript variants that fall into the previous category but have transcript models that are conserved in lesser than three of the tested species). Interestingly, all five SP prediction software that we employed predicted the presence of a SP in the amino terminus of ROR2-v1 but absence in ROR2-v2 (Table 2 and Fig. 1D).

#### Analysis of ROR1 gene and transcript variant expression in healthy human tissues

ROR1 mRNA expression for all transcripts combined and for individual transcripts was downloaded from the Genotype-Tissue Expression (GTEx) portal. Low ROR1 mRNA expression (<10 TPM) was found in most adult human tissues (Fig. 2A). Tissues with the highest ROR1 expression in the human body included arterial and cervical tissues (Fig. 2A). Surprisingly, although ROR1-v1 was predicted as the functional isoform for ROR1 (Table 1), analysis of mRNA expression of transcript variants of ROR1 identified ROR1-v3 (the variant lacking the SP sequence) as the most highly expressed isoform in adult tissues (Fig. 2B).

**Figure 2.**
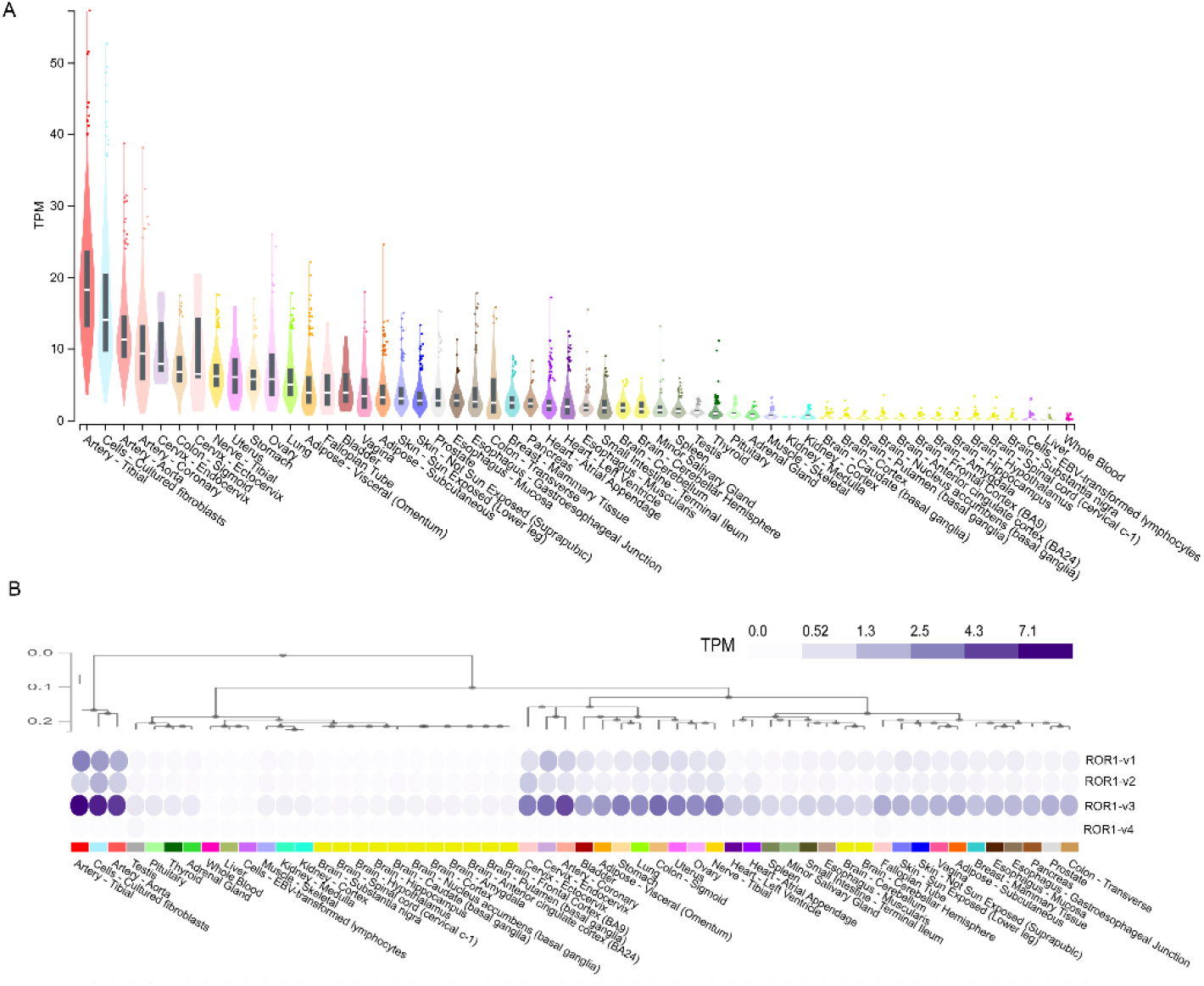
Analysis of ROR1 gene and transcript variant expression in the human body: Gene expression (A) and transcript variant expression (B) of ROR1 in 17,382 samples across 54 non-diseased human tissues from nearly 1000 individuals profiled by the GTEx project.

#### Analysis of ROR2 gene and transcript variant expression in healthy human tissues

ROR2 mRNA expression was moderately higher than ROR1 mRNA expression in adult human tissues (Fig. 3A). ROR2 mRNA expression was the highest in the sigmoid colon, esophagus and reproductive tissues including cervix, ovary, prostate, fallopian tube and vagina (Fig. 3A). Although transcript variants ROR2-v1 and ROR2-v2 were both predicted as functional isoforms of ROR2 (Table 1), analysis of mRNA expression of transcript variants of ROR2 in adult tissues identified ROR2-v1 as the most highly expressed isoform (Fig. 3B).

**Figure 3.**
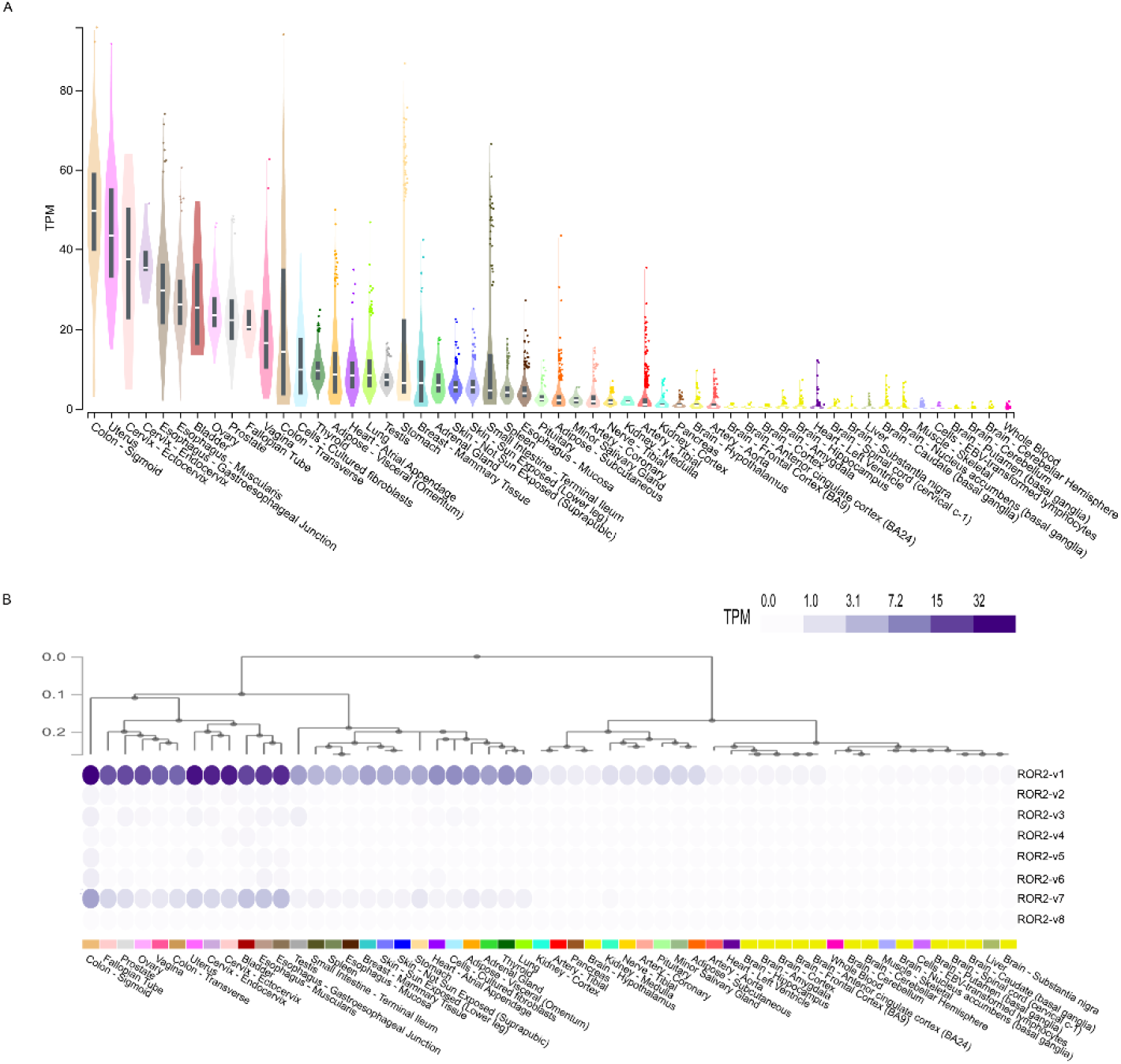
Analysis of ROR2 gene and transcript variant expression in the human body: Gene expression (A) and transcript variant expression (B) of ROR2 in 17,382 samples across 54 non-diseased human tissues from nearly 1000 individuals profiled by the GTEx project.

#### Analysis of ROR1 and ROR2 transcript variant expression in human tumour tissues

ROR1 and ROR2 transcript variant expression was analysed in 33 different tumour types profiled by TCGA (Fig. 4A and 4B). Similar to non-diseased tissues, ROR1-v3 rather than ROR1-v1 was the most highly expressed transcript variant in all tumour types analysed (Fig. 4A.). In stomach adenocarcinoma, transcript variants ROR1-v3 and ROR1-v1 were equally expressed.

**Figure 4:**
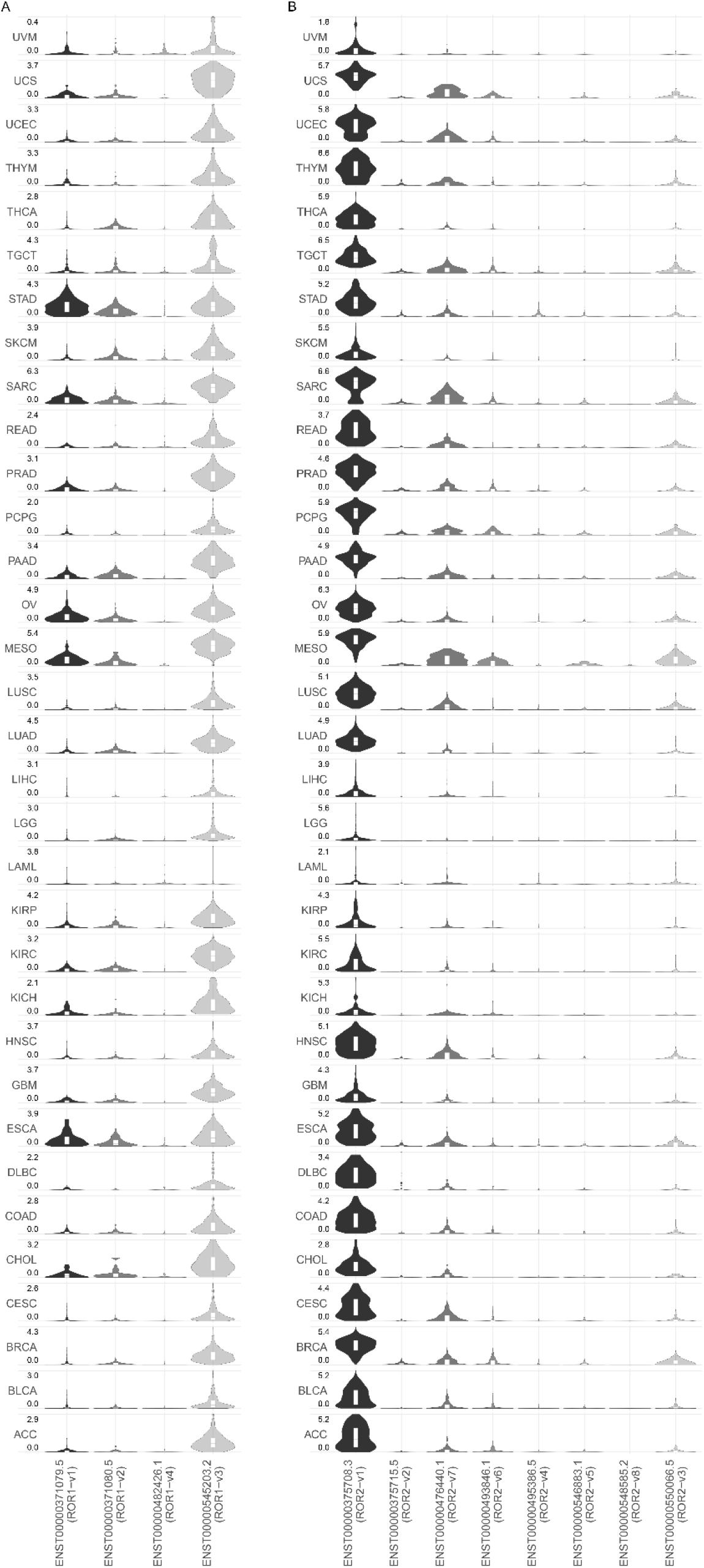
Analysis of ROR1 and ROR2 transcript variant expression in human tumour tissues: **A)** Expression of transcript variants of ROR1 in 33 tumour types examined by TCGA. B) Expression of transcript variants of ROR2 in 33 tumour types examined by TCGA. ACC (adrenocortical carcinoma), BLCA (bladder urothelial carcinoma), BRCA (breast invasive carcinoma), CESC (cervical squamous cell carcinoma and endocervical adenocarcinoma), CHOL (cholangiocarcinoma), COAD (colon adenocarcinoma), DLBC (lymphoid neoplasm diffuse large B-cell lymphoma), ESCA (esophageal carcinoma), GBM (glioblastoma multiforme), HNSC (head and neck squamous cell carcinoma), KICH (kidney chromophobe), KIRC (kidney renal clear cell carcinoma), KIRP (kidney renal papillary cell carcinoma), LAML (acute myeloid leukemia), LGG (brain lower grade glioma), LIHC (liver hepatocellular carcinoma), LUAD (lung adenocarcinoma), LUSC (lung squamous cell carcinoma), MESO (mesothelioma), OV (ovarian serous cystadenocarcinoma), PAAD (pancreatic adenocarcinoma), PCPG (pheochromocytoma and paraganglioma), PRAD (prostate adenocarcinoma), READ (rectum adenocarcinoma), SARC (sarcoma), SKCM (skin cutaneous melanoma), STAD (stomach adenocarcinoma), TGCT (testicular germ cell tumours), THCA (thyroid carcinoma), THYM (thymoma), UCEC (uterine corpus endometrial carcinoma), UCS (uterine carcinosarcoma) and UVM (uveal melanoma).

For ROR2, **s**imilar to non-diseased tissues, ROR2-v1 was the most highly expressed isoform in all 33 different tumour types profiled by TCGA (Fig. 4B). ROR2-v2, the alternate isoform predicted to be functionally important had minimal to no expression in tumour tissues.

## Discussion

ROR1 is considered an ideal druggable target for oncology because of demonstrated pro-tumourigenic actions, cancer specific expression, cell surface expression and availability of drugs that can regulate ROR1 action. Stage I clinical trials reported a ROR1 targeted monoclonal antibody, cirmtuzumab, to be well tolerated with little to no adverse effects reported [28], resulting in considerable attention being paid to this drug target. In fact, two separate ROR1-ADCs were recently acquired by Merck and Boehringer Ingelheim for multi-billion dollar amounts. ROR2 is also under preclinical evaluation as a drug target, with a number of clinical trials in progress including one of Bioatla BA3021, a conditionally active biologic (CAB) ROR2-targeted ADC (https://clinicaltrials.gov/ct2/show/NCT03504488).

Contrary to early reports that ROR1 was not expressed in adult tissues, studies have found ROR1 expression in several healthy adult tissues, highlighting possible on-target off-tumour effects of targeted therapy [16]. Balakrishnan et al. reported that a novel ROR1 antibody (6D4) identified ROR1 protein expression in several healthy human tissues [16]. RNAseq can be substantially more sensitive than IHC and immunoblotting, the commonly used techniques for examining protein expression [29, 30]. We examined ROR1 and ROR2 mRNA expression in the GTEx RNAseq transcriptomic dataset to establish expression of the pathway in an extensive array of non-diseased human tissues. As shown in Fig. 2A and 3A, several adult human tissues did express low levels of ROR1 and low to medium levels of ROR2 mRNA; hence, expression was not completely absent as suggested in early studies [5, 31, 32]. Importantly, arterial tissues expressed the highest levels of ROR1 mRNA (Fig. 2A), highlighting a need to monitor on-target off-tumour effects of ROR1 therapy on the circulatory system. Highest ROR2 expression was in the sigmoid colon, esophagus and reproductive tissues (Fig. 3A). Cultured fibroblasts have amongst the highest levels of both ROR1/2 expression (Fig. 2A and 3A). This is important as cultured fibroblasts are increasingly used in 3D or co-culture models of metastasis, and active Wnt signalling could alter cell behaviour in downstream assays. It is important to note that the activity of a pathway is controlled at multiple points including regulation of mRNA and protein expression, stability of mRNA transcript and protein, post-translational modifications, amongst others. The mere presence of transcripts does not guarantee that the pathway is functional. Conversely, gene expression can be maintained at low basal levels and dramatically upregulated upon stimulation. Hence, low expression of ROR1 and ROR2 does not rule out the possibility that these genes are further upregulated under certain conditions. Furthermore, the mRNA expression data examined in this paper covers a large array of human tissues; however, the data represents expression in bulk tissues comprising of various cell types and hence does not rule out possible higher expression in specific cell types with underrepresented cell numbers in the tissue.

ROR1 and ROR2 are reported as cell surface receptors activated by Wnt5a. The ROR1 targeted therapeutic strategies that have reached clinical trials include a monoclonal antibody and CAR-T cells targeting ROR1 [11, 28]. These agents are primarily designed for a cell surface protein; however, multiple studies have found cytoplasmic rather than plasma membrane staining for ROR1 [10, 12, 16]. Furthermore, if ROR1 is not localised to the cell surface, it theoretically cannot bind Wnt5A, the currently accepted mechanism for ROR1 mediated tumourigenesis [5]. Conflicting reports regarding the pro-tumourigenic roles of ROR2 have been reported, with studies showing both higher and lower expression to be associated with poor survival [6, 17, 20, 21]. A possible reason for these inconsistencies in localisation and/or function between studies could be the choice of the gene isoform that was examined in each study. While ROR1 and ROR2 have several isoforms, the functional isoform/s for each gene has not previously been systematically established. This study first established the various isoforms identified in the GRCh38 human reference genome (Table 1), followed by examining the expression pattern of these isoforms in both non-diseased and cancerous human tissues (Fig. 2–4), and possible functional difference between the most highly expressed isoform and the previously predicted functional isoform (Fig. 1).

ROR1-v1 was predicted as the functional isoform by the annotation pipelines APPRIS and MANE (Table 1). The TSL method examines if a transcript model (the specific exons that makeup a transcript) predicted by GENCODE is supported by mRNA expression sequences from the International Nucleotide Sequence Database Collaboration (INSDC). ROR1-v3 has a TSL5 rating which means that no single transcript from data available with the INSDC supports the GENCODE model structure for this variant. However, GTEx as well as the 33 TCGA datasets report the ROR1-v3 variant. Furthermore, surprisingly ROR1-v3 appears to be the predominantly expressed variant in both non-diseased and tumour tissues (Fig. 2B and 4). ROR1-v3 lacks a SP that ROR1-v1 contains for cell surface localisation (Fig. 1B and Table 2). This small difference could have profound implications: the protein product of ROR1-v3 may be expressed within the cell and is a possible explanation for numerous studies employing different antibodies reporting cytoplasmic staining for ROR1 [7, 10, 12–15]. ROR1-v2 was presumed to be responsible for the cytoplasmic localisation [16]; however, our analysis demonstrates that this variant also has a SP for membrane expression (Fig. 1B and Table 2). Importantly, the current immunotherapy targeting a cell surface ROR1 would not affect a cytoplasmic isoform. Given that cytoplasmic ROR1 actions would be expected to be independent of Wnt5A mediated membrane ROR1 signalling, the role of this isoform remains to be explored. In light of the ROR1-v3 variant expression, experimental strategies need to be taken into careful consideration when interpreting the role of ROR1; for e.g., overexpression constructs of ROR1 would represent only one of the variants, while knockdown probes, antibodies and primers would most likely target both of the above ROR1 variants. Hence it is possible that expression studies employing antibodies and primers may have actually analysed the ROR1-v3 variant but functional studies employing expression vectors may have analysed the ROR1-v1 variant. Further studies monitoring the two variants of ROR1 with a c-terminal tag can confirm the difference in subcellular localisation of the two variants. The start site of the ROR1-v3 variant also requires experimental confirmation.

If the argument that the non-principal variant (ROR1-v3) is not functionally important hold true, then it is important to note that analyses of expression quantitation that employ antibodies, nucleotide primers and probes that recognise both variants may not be able to differentiate between the two variants and may be confounded by the highly expressed ROR1-v3 variant. The likelihood of this would appear to be high because the difference between the two variants is merely the first exon.

Of the eight isoforms of ROR2 identified on Ensemble (release 104), ROR2-v1 is predicted as principal isoforms by APPRIS (Table 1). Interestingly, APPRIS annotation on previous releases of Ensemble predicted both ROR2-v1 and ROR2-v2 as principal isoforms (Table 1). Similar to ROR1, our analysis demonstrates a difference in predicted sub-cellular localisation between the two variants, with the presence and absence of a SP in ROR2-v1 and ROR2-v2 respectively (Fig. 1D and Table 2). The predicted cell surface localisation of ROR2-v1 and cytoplasmic localisation of ROR2-v2 could result in profound functional differences between the two variants. ROR2-v1 was the predominantly expressed isoform in both non-diseased tissues and among all 33 different tumour types profiled by TCGA (Fig. 3B and 4B) and hence this is the isoform that should primarily be examined when delineating the role of ROR2 in cancer. Expression analyses and functional studies involving ROR2-v2 alone could result in erroneous interpretation of results.

Early studies reported the proteins to possess kinase activity [33–35]; however, these findings were later disputed by studies that identified the absence of conserved residues essential for phosphotransferase activity within the tyrosine kinase-like domains of ROR1 and ROR2 [36, 37]. While RORs are now mostly accepted as pseudo-kinases, irrefutable evidence for this remains to be found. We sought to examine if there were differences related to the kinase domains between the transcript variants. ROR1-v1 and ROR1-v3 contained the kinase domains while it was absent in ROR1-v2 (Fig. 1B). For ROR2, the kinase domains were identical between both the v1 and v2 protein coding transcript variants (Fig. 1D).

### Conclusions

This study confirms mRNA expression of both ROR1 and ROR2 (albeit low levels) in non-diseased human tissues and hence, on-target off-tumour effects of targeted therapy need to be closely monitored. Surprisingly, the most highly expressed isoform of ROR1 in both non-diseased and tumour tissues lacks a SP for cell surface localisation and hence may be expressed within the cell, therefore highlighting several important questions regarding yet unknown functions of this isoform and efficacy of therapies targeting primarily a cell surface isoform. Differences in sub-cellular localisation between the transcript variants of ROR1 and ROR2 highlight a need for careful interpretation of past studies.

## Methods

### Sequence analysis of transcript variants of human ROR1 and ROR2 genes

Transcript variants of ROR1 and ROR2 identified in the GRCh38 human reference genome assembly were assessed on the Ensemble browser (release 104) on 2-Aug-2021. Results from GENCODE, Transcript Support Level, APPRIS and MANE Select annotation pipelines employed to identify functionally relevant transcript variants were examined for both genes. The protein sequences of the coding variants of ROR1 and ROR2 were analysed with multiple SP prediction software (SignalP 5.0, DeepSig, Philius, Phobius and Predisi) to identify cell surface localisation.

### Analysis of ROR1 and ROR2 gene and transcript variant expression in healthy human tissues

ROR1 and ROR2 mRNA expression data in 17,382 samples across 54 non-diseased human tissues from nearly 1000 individuals was downloaded from the GTEx portal on 10-Mar-2020. Expression of transcript variants of ROR1 and ROR2 were downloaded from the GTEx portal on 10-Mar-2020.

### Analysis of ROR1 and ROR2 transcript variant expression in human tumour tissues

Expression of transcript variants of ROR1 and ROR2 in 33 tumour types examined by TCGA was assessed on the Gene Expression Profiling Interactive Analysis (GEPIA) 2 server on 16-Aug-2021 [38].

## Acknowledgements

The results presented here are in whole based upon data generated by the TCGA Research Network: https://www.cancer.gov/tcga and the GTEx project (https://www.cancer.gov/tcga). The GTEx project was supported by the Common Fund of the Office of the Director of the National Institutes of Health, and by NCI, NHGRI, NHLBI, NIDA, NIMH, and NINDS. We gratefully acknowledge the contribution of all specimen donors and research groups involved the TCGA and GTEx projects.

## Author contributions

**Miya John:** Conceptualization, Methodology, Investigation, Data Curation, Writing - Original draft, review and editing, Visualization, Funding acquisition

**Caroline Ford:** Conceptualization, Resources, Writing - review and editing, Supervision, Project administration, Funding acquisition

