## Supplementary Figure 1 for "Pan-tissue and -cancer analysis of ROR1 and ROR2 transcript variants identify novel functional significance for an alternative splice variant of ROR1"

**S1. cDNA sequence for ROR1 transcripts:** Multiple sequence alignment of full-length cDNA sequence of ROR1 transcript variants ROR1-v1 (ENST00000371079.6), ROR1-v2 (ENST00000371080.5) and ROR1-v3 (ENST00000545203.2). Alternating exons are represented in black and blue. Coding sequence is represented in bold text. Start codons of ROR1-v1 and ROR1-v2 are represented in red. The first five ATG codons of ROR1-v3 are highlighted in yellow and alternating codons are underscored.

```
ROR1-v2      -----GAGCGAGAGAGGGAGCGTGGAGAGCTGGAGCAGCCGCCACCGCCGCCGCCGAGGG	55
ROR1-v1      AAGTTGAGCGAGAGAGGGAGCGTGGAGAGCTGGAGCAGCCGCCACCGCCGCCGCCGAGGG	60
ROR1-v3      ------------------------------------------------------------	0


ROR1-v2      AGCCCCGGGACGGCAGCCCCTGGGCGCAGGGTGCGCTGTTCTCGGAGTCCGACCCAGGGC	115
ROR1-v1      AGCCCCGGGACGGCAGCCCCTGGGCGCAGGGTGCGCTGTTCTCGGAGTCCGACCCAGGGC	120
ROR1-v3      ------------------------------------------------------------	0


ROR1-v2      GACTCACGCCCACTGGTGCGACCCGGACAGCCTGGGACTGACCCGCCGGCCCAGGCGAGG	175
ROR1-v1      GACTCACGCCCACTGGTGCGACCCGGACAGCCTGGGACTGACCCGCCGGCCCAGGCGAGG	180
ROR1-v3      ------------------------------------------------------------	0


ROR1-v2      CTGCAGCCAGAGGGCTGGGAAGGGATCGCGCTCGCGGCATCCAGAGGCGGCCAGGCGGAG	235
ROR1-v1      CTGCAGCCAGAGGGCTGGGAAGGGATCGCGCTCGCGGCATCCAGAGGCGGCCAGGCGGAG	240
ROR1-v3      ------------------------------------------------------------	0


ROR1-v2      GCGAGGGAGCAGGTTAGAGGGACAAAGAGCTTTGCAGACGTCCCCGGCGTCCTGCGAGCG	295
ROR1-v1      GCGAGGGAGCAGGTTAGAGGGACAAAGAGCTTTGCAGACGTCCCCGGCGTCCTGCGAGCG	300
ROR1-v3      ------------------------------------------------------------	0


ROR1-v2      CCAGCGGCCGGGACGAGGCGGCCGGGAGCCCGGGAAGAGCCCGTGGATGTTCTGCGCGCG	355
ROR1-v1      CCAGCGGCCGGGACGAGGCGGCCGGGAGCCCGGGAAGAGCCCGTGGATGTTCTGCGCGCG	360
ROR1-v3      ------------------------------------------------------------	0


ROR1-v2      GCCTGGGAGCCGCCGCCGCCGCCGCCTCAGCGAGAGGAGGAATGCACCGGCCGCGCCGCC	415
ROR1-v1      GCCTGGGAGCCGCCGCCGCCGCCGCCTCAGCGAGAGGAGGAATGCACCGGCCGCGCCGCC	420
ROR1-v3      ------------------------------------------------------------	0


ROR1-v2      GCGGGACGCGCCCGCCGCTCCTGGCGCTGCTGGCCGCGCTGCTGCTGGCCGCACGCGGGG	475
ROR1-v1      GCGGGACGCGCCCGCCGCTCCTGGCGCTGCTGGCCGCGCTGCTGCTGGCCGCACGCGGGG	480
ROR1-v3      ------------------------------------------------------------	0


ROR1-v2      CTGCTGCCCAAGAAACAGAGCTGTCAGTCAGTGCTGAATTAGTGCCTACCTCATCATGGA	535
ROR1-v1      CTGCTGCCCAAGAAACAGAGCTGTCAGTCAGTGCTGAATTAGTGCCTACCTCATCATGGA	540
ROR1-v3      ----------AGAAACAGAGCTGTCAGTCAGTGCTGAATTAGTGCCTACCTCATCATGGA	50
                       **************************************************

ROR1-v2      ACATCTCAAGTGAACTCAACAAAGATTCTTACCTGACCCTCGATGAACCAATGAATAACA	595
ROR1-v1      ACATCTCAAGTGAACTCAACAAAGATTCTTACCTGACCCTCGATGAACCAATGAATAACA	600
ROR1-v3      ACATCTCAAGTGAACTCAACAAAGATTCTTACCTGACCCTCGATGAACCAATGAATAACA	110
             ************************************************************

ROR1-v2      TCACCACGTCTCTGGGCCAGACAGCAGAACTGCACTGCAAAGTCTCTGGGAATCCACCTC	655
ROR1-v1      TCACCACGTCTCTGGGCCAGACAGCAGAACTGCACTGCAAAGTCTCTGGGAATCCACCTC	660
ROR1-v3      TCACCACGTCTCTGGGCCAGACAGCAGAACTGCACTGCAAAGTCTCTGGGAATCCACCTC	170
             ************************************************************

ROR1-v2      CCACCATCCGCTGGTTCAAAAATGATGCTCCTGTGGTCCAGGAGCCCCGGAGGCTCTCCT	715
ROR1-v1      CCACCATCCGCTGGTTCAAAAATGATGCTCCTGTGGTCCAGGAGCCCCGGAGGCTCTCCT	720
ROR1-v3      CCACCATCCGCTGGTTCAAAAATGATGCTCCTGTGGTCCAGGAGCCCCGGAGGCTCTCCT	230
             ************************************************************

ROR1-v2      TTCGGTCCACCATCTATGGCTCTCGGCTGCGGATTAGAAACCTCGACACCACAGACACAG	775
ROR1-v1      TTCGGTCCACCATCTATGGCTCTCGGCTGCGGATTAGAAACCTCGACACCACAGACACAG	780
ROR1-v3      TTCGGTCCACCATCTATGGCTCTCGGCTGCGGATTAGAAACCTCGACACCACAGACACAG	290
             ************************************************************

ROR1-v2      GCTACTTCCAGTGCGTGGCAACAAACGGCAAGGAGGTGGTTTCTTCCACTGGAGTCTTGT	835
ROR1-v1      GCTACTTCCAGTGCGTGGCAACAAACGGCAAGGAGGTGGTTTCTTCCACTGGAGTCTTGT	840
ROR1-v3      GCTACTTCCAGTGCGTGGCAACAAACGGCAAGGAGGTGGTTTCTTCCACTGGAGTCTTGT	350
             ************************************************************

ROR1-v2      TTGTCAAGTTTGGCCCCCCTCCCACTGCAAGTCCAGGATACTCAGATGAGTATGAAGAAG	895
ROR1-v1      TTGTCAAGTTTGGCCCCCCTCCCACTGCAAGTCCAGGATACTCAGATGAGTATGAAGAAG	900
ROR1-v3      TTGTCAAGTTTGGCCCCCCTCCCACTGCAAGTCCAGGATACTCAGATGAGTATGAAGAAG	410
             ************************************************************

ROR1-v2      ATGGATTCTGTCAGCCATACAGAGGGATTGCATGTGCAAGATTTATTGGCAACCGCACCG	955
ROR1-v1      ATGGATTCTGTCAGCCATACAGAGGGATTGCATGTGCAAGATTTATTGGCAACCGCACCG	960
ROR1-v3      ATGGATTCTGTCAGCCATACAGAGGGATTGCATGTGCAAGATTTATTGGCAACCGCACCG	470
             ************************************************************

ROR1-v2      TCTATATGGAGTCTTTGCACATGCAAGGGGAAATAGAAAATCAGATCACAGCTGCCTTCA	1015
ROR1-v1      TCTATATGGAGTCTTTGCACATGCAAGGGGAAATAGAAAATCAGATCACAGCTGCCTTCA	1020
ROR1-v3      TCTATATGGAGTCTTTGCACATGCAAGGGGAAATAGAAAATCAGATCACAGCTGCCTTCA	530
             ************************************************************

ROR1-v2      CTATGATTGGCACTTCCAGTCACTTATCTGATAAGTGTTCTCAGTTCGCCATTCCTTCCC	1075
ROR1-v1      CTATGATTGGCACTTCCAGTCACTTATCTGATAAGTGTTCTCAGTTCGCCATTCCTTCCC	1080
ROR1-v3      CTATGATTGGCACTTCCAGTCACTTATCTGATAAGTGTTCTCAGTTCGCCATTCCTTCCC	590
             ************************************************************

ROR1-v2      TGTGCCACTATGCCTTCCCGTACTGCGATGAAACTTCATCCGTCCCAAAGCCCCGTGACT	1135
ROR1-v1      TGTGCCACTATGCCTTCCCGTACTGCGATGAAACTTCATCCGTCCCAAAGCCCCGTGACT	1140
ROR1-v3      TGTGCCACTATGCCTTCCCGTACTGCGATGAAACTTCATCCGTCCCAAAGCCCCGTGACT	650
             ************************************************************

ROR1-v2      TGTGTCGCGATGAATGTGAAATCCTGGAGAATGTCCTGTGTCAAACAGAGTACATTTTTG	1195
ROR1-v1      TGTGTCGCGATGAATGTGAAATCCTGGAGAATGTCCTGTGTCAAACAGAGTACATTTTTG	1200
ROR1-v3      TGTGTCGCGATGAATGTGAAATCCTGGAGAATGTCCTGTGTCAAACAGAGTACATTTTTG	710
             ************************************************************

ROR1-v2      CAAGATCAAATCCCATGATTCTGATGAGGCTGAAACTGCCAAACTGTGAAGATCTCCCCC	1255
ROR1-v1      CAAGATCAAATCCCATGATTCTGATGAGGCTGAAACTGCCAAACTGTGAAGATCTCCCCC	1260
ROR1-v3      CAAGATCAAATCCCATGATTCTGATGAGGCTGAAACTGCCAAACTGTGAAGATCTCCCCC	770
             ************************************************************

ROR1-v2      AGCCAGAGAGCCCAGAAGCTGCGAACTGTATCCGGATTGGAATTCCCATGGCAGATCCTA	1315
ROR1-v1      AGCCAGAGAGCCCAGAAGCTGCGAACTGTATCCGGATTGGAATTCCCATGGCAGATCCTA	1320
ROR1-v3      AGCCAGAGAGCCCAGAAGCTGCGAACTGTATCCGGATTGGAATTCCCATGGCAGATCCTA	830
             ************************************************************

ROR1-v2      TAAATAAAAATCACAAGTGTTATAACAGCACAGGTGTGGACTACCGGGGGACCGTCAGTG	1375
ROR1-v1      TAAATAAAAATCACAAGTGTTATAACAGCACAGGTGTGGACTACCGGGGGACCGTCAGTG	1380
ROR1-v3      TAAATAAAAATCACAAGTGTTATAACAGCACAGGTGTGGACTACCGGGGGACCGTCAGTG	890
             ************************************************************

ROR1-v2      TGACCAAATCAGGGCGCCAGTGCCAGCCATGGAATTCCCAGTATCCCCACACACACACTT	1435
ROR1-v1      TGACCAAATCAGGGCGCCAGTGCCAGCCATGGAATTCCCAGTATCCCCACACACACACTT	1440
ROR1-v3      TGACCAAATCAGGGCGCCAGTGCCAGCCATGGAATTCCCAGTATCCCCACACACACACTT	950
             ************************************************************

ROR1-v2      TCACCGCCCTTCGTTTCCCAGAGCTGAATGGAGGCCATTCCTACTGCCGCAACCCAGGGA	1495
ROR1-v1      TCACCGCCCTTCGTTTCCCAGAGCTGAATGGAGGCCATTCCTACTGCCGCAACCCAGGGA	1500
ROR1-v3      TCACCGCCCTTCGTTTCCCAGAGCTGAATGGAGGCCATTCCTACTGCCGCAACCCAGGGA	1010
             ************************************************************

ROR1-v2      ATCAAAAGGAAGCTCCCTGGTGCTTCACCTTGGATGAAAACTTTAAGTCTGATCTGTGTG	1555
ROR1-v1      ATCAAAAGGAAGCTCCCTGGTGCTTCACCTTGGATGAAAACTTTAAGTCTGATCTGTGTG	1560
ROR1-v3      ATCAAAAGGAAGCTCCCTGGTGCTTCACCTTGGATGAAAACTTTAAGTCTGATCTGTGTG	1070
             ************************************************************

ROR1-v2      ACATCCCAGCGTGCGGTAAATAGAAGTCATTG----------------------------	1587
ROR1-v1      ACATCCCAGCGTGCGATTCAAAGGATTCCAAGGAGAAGAATAAAATGGAAATCCTGTACA	1620
ROR1-v3      ACATCCCAGCGTGCGATTCAAAGGATTCCAAGGAGAAGAATAAAATGGAAATCCTGTACA	1130
             *************** *  * ** * **   *

ROR1-v2      ------CCCCTAATGTATTCAATCATCTTTAAAGATCCCTATCCTACCCCTCTTATTTA-	1640
ROR1-v1      TACTAGTGCCAAGTGTGGCCATTCCCCTGG--CCATTGCTT---TACTCTTCTTCTTCAT	1675
ROR1-v3      TACTAGTGCCAAGTGTGGCCATTCCCCTGG--CCATTGCTT---TACTCTTCTTCTTCAT	1185
                     ** * ***   ** **  **      **  **    *** * **** ** *

ROR1-v2      -----------GGA-------GAATCC-T----------ATAAGGGGGGCAAAGA-----	1666
ROR1-v1      TTGCGTCTGTCGGAATAACCAGAAGTCATCGTCGGCACCAGTCCAGAGGCAACCAAAACA	1735
ROR1-v3      TTGCGTCTGTCGGAATAACCAGAAGTCATCGTCGGCACCAGTCCAGAGGCAACCAAAACA	1245
                        ***       ***  * *          *     * *****  *

ROR1-v2      ------------AAATGGACAGT---ATTTGCTTG----AT-------------------	1688
ROR1-v1      CGTCAGAGGTCAAAATGTAGAGATGTCAATGCTGAATGCATATAAACCCAAGAGCAAGGC	1795
ROR1-v3      CGTCAGAGGTCAAAATGTAGAGATGTCAATGCTGAATGCATATAAACCCAAGAGCAAGGC	1305
                         ***** * **       ****      **

ROR1-v2      -----------CTCAATCTG------GTTTTAGGGTAAACCTT---------GCCGTTTC	1722
ROR1-v1      TAAAGAGCTACCTCTTTCTGCTGTACGCTTTATGGAAGAATTGGGTGAGTGTGCCTTTGG	1855
ROR1-v3      TAAAGAGCTACCTCTTTCTGCTGTACGCTTTATGGAAGAATTGGGTGAGTGTGCCTTTGG	1365
                        ***  ****      * **** ** * *  *          *** **

ROR1-v2      -----TACATAAAACACCTCGTA-----------AGGTACCAA----AACA---CGTTCT	1759
ROR1-v1      AAAAATCTATAAAGGCCATCTCTATCTCCCAGGCATGGACCATGCTCAGCTGGTTGCTAT	1915
ROR1-v3      AAAAATCTATAAAGGCCATCTCTATCTCCCAGGCATGGACCATGCTCAGCTGGTTGCTAT	1425
                  *  *****   * **              * * ****     * *     * * *

ROR1-v2      CAAGAAGTCAACTGCCTTTA--TACCTGCAGCCATTGCACTCATGGATGTAACAGGGACC	1817
ROR1-v1      CAAGACCTTGAAAGACTATAACAACCCCCAGCAATGGACGGAAT---TTCAACAAGAAGC	1972
ROR1-v3      CAAGACCTTGAAAGACTATAACAACCCCCAGCAATGGACGGAAT---TTCAACAAGAAGC	1482
             *****  *  *  * ** **   ***  **** ** *     **   *  **** * * *

ROR1-v2      C----------------------AGCC-------------CTT-CAG------------A	1829
ROR1-v1      CTCCCTAATGGCAGAACTGCACCACCCCAATATTGTCTGCCTTCTAGGTGCCGTCACTCA	2032
ROR1-v3      CTCCCTAATGGCAGAACTGCACCACCCCAATATTGTCTGCCTTCTAGGTGCCGTCACTCA	1542
             *                      * **             ***  **            *

ROR1-v2      GGCACA----------------GTTGAGACAGTTTATCACATTGATTTTTAT--------	1865
ROR1-v1      GGAACAACCTGTGTGCATGCTTTTTGAGTATATTAATCAGGGGGATCTCCATGAGTTCCT	2092
ROR1-v3      GGAACAACCTGTGTGCATGCTTTTTGAGTATATTAATCAGGGGGATCTCCATGAGTTCCT	1602
             ** ***                 *****    ** ****    *** *  **

ROR1-v2      ---------------AGAAAAAGATGTTAC-------------CCAGAATGGTCTGC---	1894
ROR1-v1      CATCATGAGATCCCCACACTCTGATGTTGGCTGCAGCAGTGATGAAGATGGGACTGTGAA	2152
ROR1-v3      CATCATGAGATCCCCACACTCTGATGTTGGCTGCAGCAGTGATGAAGATGGGACTGTGAA	1662
                            * *    ******                 ***  ** ***

ROR1-v2      ----------GTCCAAGTGGACCTTTTCAGCAAA--AAAAGGAATATTGGAAGCAGGAAG	1942
ROR1-v1      ATCCAGCCTGGACCA--CGGAGATTTTCTGCACATTGCAATTCAGATTGCAGCTGGCATG	2210
ROR1-v3      ATCCAGCCTGGACCA--CGGAGATTTTCTGCACATTGCAATTCAGATTGCAGCTGGCATG	1720
                       * ***   ***  ***** *** *    **   * **** *    * * *

ROR1-v2      AAATTGTTTTCT---GTATGCCTTAAGAACACCACAAGGCAGGATGAATCTACAAC--CA	1997
ROR1-v1      GAATACCTGTCTAGTCACTTCTTTG-----TCCACAAGGA-CCTTGCAGCTCGCAATATT	2264
ROR1-v3      GAATACCTGTCTAGTCACTTCTTTG-----TCCACAAGGA-CCTTGCAGCTCGCAATATT	1774
              ***   * ***      * * **       ********     ** * **   *

ROR1-v2      TTACTCGGTCA----------------------------------TCCAGGACAATCTG-	2022
ROR1-v1      TTAATCGGAGAGCAACTTCATGTAAAGATTTCAGACTTGGGGCTTTCCAGAGAAATTTAC	2324
ROR1-v3      TTAATCGGAGAGCAACTTCATGTAAAGATTTCAGACTTGGGGCTTTCCAGAGAAATTTAC	1834
             *** ****  *                                  *****   *** *

ROR1-v2      ----TGGGT-A-AACTGTGTCCTTCGT----TATGTCTGTTAATACTGCAGAAGAAGCAT	2072
ROR1-v1      TCCGCTGATTACTACAGGGTCCAGAGTAAGTCCTTGCTGCCCATTCGCTGGATGCCCCCT	2384
ROR1-v3      TCCGCTGATTACTACAGGGTCCAGAGTAAGTCCTTGCTGCCCATTCGCTGGATGCCCCCT	1894
                   * * *  ** * ****   **      *  ***   ** *    ** *   * *

ROR1-v2      ATAGGTATCTAGTAAGAAAATGGAATTCCTGAGTCAGTTAACTGTTCCTTTT--------	2124
ROR1-v1      GAAGCCATCATGTATGGCAAATTCTCTTCTGATTCAGATATCTGGTCCTTTGGGGTTGTC	2444
ROR1-v3      GAAGCCATCATGTATGGCAAATTCTCTTCTGATTCAGATATCTGGTCCTTTGGGGTTGTC	1954
               **  ***  *** *  **      * **** **** ** *** ******

ROR1-v2      -TCTAGAAAATGT------TTGGA-----------------------------GAAAATA	2148
ROR1-v1      TTGTGGGAGATTTTCAGTTTTGGACTCCAGCCATATTATGGATTCAGTAACCAGGAAGTG	2504
ROR1-v3      TTGTGGGAGATTTTCAGTTTTGGACTCCAGCCATATTATGGATTCAGTAACCAGGAAGTG	2014
              * * * * ** *      *****                             * ** *

ROR1-v2      ATGAAAATGGGCCAAGCATGGTGGCTTATACCTGTAATCCCAACACT-------------	2195
ROR1-v1      ATTGAGATGGTGAGAAAACGGCAGCTCTTACCA-TGCTCTGAAGACTGCCCACCCAGAAT	2563
ROR1-v3      ATTGAGATGGTGAGAAAACGGCAGCTCTTACCA-TGCTCTGAAGACTGCCCACCCAGAAT	2073
             **  * ****    *  * **  ***  ****  *  **  ** ***

ROR1-v2      CTA---------GGAAG-GCCGAGGCAGGAGGATCATT---TGAGCCCAGGGGTTCAAGA	2242
ROR1-v1      GTACAGCCTCATGACAGAGTGCTGGAATGAGATTCCTTCTAGGAGACCAAGATTTAAAGA	2623
ROR1-v3      GTACAGCCTCATGACAGAGTGCTGGAATGAGATTCCTTCTAGGAGACCAAGATTTAAAGA	2133
              **         *  ** *    ** * ***  ** **    *** *** *  ** ****

ROR1-v2      C---------CAGCCCAGG----------CAACATAGTGAGACTCCATCTCTACCAAAA-	2282
ROR1-v1      TATTCACGTCCGGCTTCGGTCCTGGGAGGGACTCTCAAGTCACACAAGCTCTACTACTCC	2683
ROR1-v3      TATTCACGTCCGGCTTCGGTCCTGGGAGGGACTCTCAAGTCACACAAGCTCTACTACTCC	2193
                       * **   **           *   *   *  ** * * ****** *

ROR1-v2      ----AAAAGAAAA--AAAAAAAAAGAAAA-------------------------------	2305
ROR1-v1      TTCAGGGGGAAATGCCACCACACAGACAACCTCCCTCAGTGCCAGCCCAGTGAGTAATCT	2743
ROR1-v3      TTCAGGGGGAAATGCCACCACACAGACAACCTCCCTCAGTGCCAGCCCAGTGAGTAATCT	2253
                     ****    *  * * *** **

ROR1-v2      ------------------------------------------------------------	2305
ROR1-v1      CAGTAACCCCAGATATCCTAATTACATGTTCCCGAGCCAGGGTATTACACCACAGGGCCA	2803
ROR1-v3      CAGTAACCCCAGATATCCTAATTACATGTTCCCGAGCCAGGGTATTACACCACAGGGCCA	2313


ROR1-v2      ------------------------------------------------------------	2305
ROR1-v1      GATTGCTGGTTTCATTGGCCCGCCAATACCTCAGAACCAGCGATTCATTCCCATCAATGG	2863
ROR1-v3      GATTGCTGGTTTCATTGGCCCGCCAATACCTCAGAACCAGCGATTCATTCCCATCAATGG	2373


ROR1-v2      ------------------------------------------------------------	2305
ROR1-v1      ATACCCAATACCTCCTGGATATGCAGCGTTTCCAGCTGCCCACTACCAGCCAACAGGTCC	2923
ROR1-v3      ATACCCAATACCTCCTGGATATGCAGCGTTTCCAGCTGCCCACTACCAGCCAACAGGTCC	2433


ROR1-v2      ------------------------------------------------------------	2305
ROR1-v1      TCCCAGAGTGATTCAGCACTGCCCACCTCCCAAGAGTCGGTCCCCAAGCAGTGCCAGTGG	2983
ROR1-v3      TCCCAGAGTGATTCAGCACTGCCCACCTCCCAAGAGTCGGTCCCCAAGCAGTGCCAGTGG	2493


ROR1-v2      ------------------------------------------------------------	2305
ROR1-v1      GTCGACTAGCACTGGCCATGTGACTAGCTTGCCCTCATCAGGATCCAATCAGGAAGCAAA	3043
ROR1-v3      GTCGACTAGCACTGGCCATGTGACTAGCTTGCCCTCATCAGGATCCAATCAGGAAGCAAA	2553


ROR1-v2      ------------------------------------------------------------	2305
ROR1-v1      TATTCCTTTACTACCACACATGTCAATTCCAAATCATCCTGGTGGAATGGGTATCACCGT	3103
ROR1-v3      TATTCCTTTACTACCACACATGTCAATTCCAAATCATCCTGGTGGAATGGGTATCACCGT	2613


ROR1-v2      ------------------------------------------------------------	2305
ROR1-v1      TTTTGGCAACAAATCTCAAAAACCCTACAAAATTGACTCAAAGCAAGCATCTTTACTAGG	3163
ROR1-v3      TTTTGGCAACAAATCTCAAAAACCCTACAAAATTGACTCAAAGCAAGCATCTTTACTAGG	2673


ROR1-v2      ------------------------------------------------------------	2305
ROR1-v1      AGACGCCAATATTCATGGACACACCGAATCTATGATTTCTGCAGAACTGTAAAATGCACA	3223
ROR1-v3      AGACGCCAATATTCATGGACACACCGAATCTATGATTTCTGCAGAACTGTAAAATGCACA	2733


ROR1-v2      ------------------------------------------------------------	2305
ROR1-v1      ACTTTTGTAAATGTGGTATACAGGACAAACTAGACGGCCGTAGAAAAGATTTATATTCAA	3283
ROR1-v3      ACTTTTGTAAATGTGGTATACAGGACAAACTAGACGGCCGTAGAAAAGATTTATATTCAA	2793


ROR1-v2      ------------------------------------------------------------	2305
ROR1-v1      ATGTTTTTATTAAAGTAAGGTTCTCATTTAGCAGACATCGCAACAAGTACCTTCTGTGAA	3343
ROR1-v3      ATGTTTTTATTAAAGTAAGGTTCTCATTTAGCAGACATCGCAACAAGTACCTTCTGTGAA	2853


ROR1-v2      ------------------------------------------------------------	2305
ROR1-v1      GTTTCACTGTGTCTTACCAAGCAGGACAGACACTCGGCCAGAAAAAAAAAAAAAAAAAAA	3403
ROR1-v3      GTTTCACTGTGTCTTACCAAGCAGGACAGACACTCGGCCAGAAAAAAAAAAAAAAAAAAA	2913


ROR1-v2      ------------------------------------------------------------	2305
ROR1-v1      AAACAAGCAAACAAAAACATTGTGGGATGTGCACTCCATTGGAGTGCATGACATGGCATT	3463
ROR1-v3      AAACAAGCAAACAAAAACATTGTGGGATGTGCACTCCATTGGAGTGCATGACATGGCATT	2973


ROR1-v2      ------------------------------------------------------------	2305
ROR1-v1      GGGATTGGAACATGTGGTTTCGAGCACTGAAAGCTGCAAACCAGTGAAGAGGAAAAGAAC	3523
ROR1-v3      GGGATTGGAACATGTGGTTTCGAGCACTGAAAGCTGCAAACCAGTGAAGAGGAAAAGAAC	3033


ROR1-v2      ------------------------------------------------------------	2305
ROR1-v1      CTTGTGATTAAATATAAAACCAAAAGTCAAATGGTGCTTTGTGTTTTAGCCTTCAGTCAC	3583
ROR1-v3      CTTGTGATTAAATATAAAACCAAAAGTCAAATGGTGCTTTGTGTTTTAGCCTTCAGTCAC	3093


ROR1-v2      ------------------------------------------------------------	2305
ROR1-v1      CATGACTGGTCTCTCCCCCAGATGTATATATACCATAGCATTTGTCTACCTGCTGTCTTT	3643
ROR1-v3      CATGACTGGTCTCTCCCCCAGATGTATATATACCATAGCATTTGTCTACCTGCTGTCTTT	3153


ROR1-v2      ------------------------------------------------------------	2305
ROR1-v1      TCTTCAGGACAGATGTTCAGGAATTATATTGATTGAATTTAGACTCTGTGCATGTTCTTA	3703
ROR1-v3      TCTTCAGGACAGATGTTCAGGAATTATATTGATTGAATTTAGACTCTGTGCATGTTCTTA	3213


ROR1-v2      ------------------------------------------------------------	2305
ROR1-v1      TGGAAATGATGTTCAGAATCCATGAAGAAACTTCAGGCCAAATTTGAAACCCTGGAGGGA	3763
ROR1-v3      TGGAAATGATGTTCAGAATCCATGAAGAAACTTCAGGCCAAATTTGAAACCCTGGAGGGA	3273


ROR1-v2      ------------------------------------------------------------	2305
ROR1-v1      AATGAGCCATAAGGGAAGTATAACAAGCCCTGAAGCCTTTTATGTCGTTGTGCTTCTTTG	3823
ROR1-v3      AATGAGCCATAAGGGAAGTATAACAAGCCCTGAAGCCTTTTATGTCGTTGTGCTTCTTTG	3333


ROR1-v2      ------------------------------------------------------------	2305
ROR1-v1      GGAAGGTGTAGAGTGTGCCTTTTTGTGAATCCTCCTCTGATCATGAGGGTCTTTCCCACA	3883
ROR1-v3      GGAAGGTGTAGAGTGTGCCTTTTTGTGAATCCTCCTCTGATCATGAGGGTCTTTCCCACA	3393


ROR1-v2      ------------------------------------------------------------	2305
ROR1-v1      GTTTCTCACAGTGTGTTTACACTGCCCTTGGAATAACACAGCGCATCAGACCATAAGAAG	3943
ROR1-v3      GTTTCTCACAGTGTGTTTACACTGCCCTTGGAATAACACAGCGCATCAGACCATAAGAAG	3453


ROR1-v2      ------------------------------------------------------------	2305
ROR1-v1      GCTAGATGTGGATGCTAGAATTGATTGTTGGTTGATAGTTCTCTTTGCTGGATTAGGAAT	4003
ROR1-v3      GCTAGATGTGGATGCTAGAATTGATTGTTGGTTGATAGTTCTCTTTGCTGGATTAGGAAT	3513


ROR1-v2      ------------------------------------------------------------	2305
ROR1-v1      GAGGCGCCAAAGGAAGCACAGCCAGGAAAATGGCCCCACAGCCTAGATCAGCATCTGTGG	4063
ROR1-v3      GAGGCGCCAAAGGAAGCACAGCCAGGAAAATGGCCCCACAGCCTAGATCAGCATCTGTGG	3573


ROR1-v2      ------------------------------------------------------------	2305
ROR1-v1      GAAAGGAAAAAGGGTTCCCATGGGGACAGCCCCCATAGGAGTTTTCTGGAACCAGTAACA	4123
ROR1-v3      GAAAGGAAAAAGGGTTCCCATGGGGACAGCCCCCATAGGAGTTTTCTGGAACCAGTAACA	3633


ROR1-v2      ------------------------------------------------------------	2305
ROR1-v1      CTGAAAAATAAGTGTGTGGCTACAGATGAGCACGCCCACCCCTTGCAACTCCCTGTTTAC	4183
ROR1-v3      CTGAAAAATAAGTGTGTGGCTACAGATGAGCACGCCCACCCCTTGCAACTCCCTGTTTAC	3693


ROR1-v2      ------------------------------------------------------------	2305
ROR1-v1      AAGTTGTCCGAGGCATTGGAGTGCTTATGGTCAATGGGCTCTAGGGAAGTAGGAAACTCC	4243
ROR1-v3      AAGTTGTCCGAGGCATTGGAGTGCTTATGGTCAATGGGCTCTAGGGAAGTAGGAAACTCC	3753


ROR1-v2      ------------------------------------------------------------	2305
ROR1-v1      ATCATGATACAAATGTCTAGTAATTTTAAAGTTTCTTTCCCTTTTTTTCTGTGCTGGAAA	4303
ROR1-v3      ATCATGATACAAATGTCTAGTAATTTTAAAGTTTCTTTCCCTTTTTTTCTGTGCTGGAAA	3813


ROR1-v2      ------------------------------------------------------------	2305
ROR1-v1      TGTTCACAGATTTGATTCCCGCCCCAAGAATTACAACAATGTATTCATACAGTCAGTTGG	4363
ROR1-v3      TGTTCACAGATTTGATTCCCGCCCCAAGAATTACAACAATGTATTCATACAGTCAGTTGG	3873


ROR1-v2      ------------------------------------------------------------	2305
ROR1-v1      AATCCAAAGGCAATTAATTCTATTTTGCAAAATATGATGGTCTTCCTAAAAAACAAGTAC	4423
ROR1-v3      AATCCAAAGGCAATTAATTCTATTTTGCAAAATATGATGGTCTTCCTAAAAAACAAGTAC	3933


ROR1-v2      ------------------------------------------------------------	2305
ROR1-v1      TGAGTTCTCATTTCAAAAGTTACCAAGAACTGAATTCTTTAAACTAGCAAACCGAAGTAA	4483
ROR1-v3      TGAGTTCTCATTTCAAAAGTTACCAAGAACTGAATTCTTTAAACTAGCAAACCGAAGTAA	3993


ROR1-v2      ------------------------------------------------------------	2305
ROR1-v1      CATGACCATTTTTGCCTTAGTGGGAGGTTTCAAATGTGCAGCTCATGGCATTTACCTGCC	4543
ROR1-v3      CATGACCATTTTTGCCTTAGTGGGAGGTTTCAAATGTGCAGCTCATGGCATTTACCTGCC	4053


ROR1-v2      ------------------------------------------------------------	2305
ROR1-v1      GACCATCTTTTGCCAAGTTTAGAATTCTTATGCGTTTCAAGTTCTATATAGAAAGTAATT	4603
ROR1-v3      GACCATCTTTTGCCAAGTTTAGAATTCTTATGCGTTTCAAGTTCTATATAGAAAGTAATT	4113


ROR1-v2      ------------------------------------------------------------	2305
ROR1-v1      TTACTTTTGATTTTTCTCTTGTTAAAAAAAAACTCCTTTATTCTAAGAACAATGTCTCAA	4663
ROR1-v3      TTACTTTTGATTTTTCTCTTGTTAAAAAAAAACTCCTTTATTCTAAGAACAATGTCTCAA	4173


ROR1-v2      ------------------------------------------------------------	2305
ROR1-v1      AGTCTCATTTTTACTTTAAAGGTATAAGAGACTTCTAAAGAGACTTACGGGATATAAAAG	4723
ROR1-v3      AGTCTCATTTTTACTTTAAAGGTATAAGAGACTTCTAAAGAGACTTACGGGATATAAAAG	4233


ROR1-v2      ------------------------------------------------------------	2305
ROR1-v1      TAATTCCTGGAAATGATATTTGATGAGGAGAATGTAAGAGAATGAAAAACACATTGATGT	4783
ROR1-v3      TAATTCCTGGAAATGATATTTGATGAGGAGAATGTAAGAGAATGAAAAACACATTGATGT	4293


ROR1-v2      ------------------------------------------------------------	2305
ROR1-v1      TTTCATTTTTAAAAAATACAACTAGCATGAGAAATCAAGAAAATATGTTTACCAAAATGC	4843
ROR1-v3      TTTCATTTTTAAAAAATACAACTAGCATGAGAAATCAAGAAAATATGTTTACCAAAATGC	4353


ROR1-v2      ------------------------------------------------------------	2305
ROR1-v1      ATTGCAATTTTCCCAAACCTGAGTCTTCAAATAACAAACATGAACTTATAGGTACTGTGA	4903
ROR1-v3      ATTGCAATTTTCCCAAACCTGAGTCTTCAAATAACAAACATGAACTTATAGGTACTGTGA	4413


ROR1-v2      ------------------------------------------------------------	2305
ROR1-v1      ACTAGAAGAATTGGTTATATCCAGATTTCTGGGAGATAAATTAAAACATATTTTTGTGGC	4963
ROR1-v3      ACTAGAAGAATTGGTTATATCCAGATTTCTGGGAGATAAATTAAAACATATTTTTGTGGC	4473


ROR1-v2      ------------------------------------------------------------	2305
ROR1-v1      ATAAATAAGCTGATTCAGAAGTTACATTTCTTAACTTTAGGTAGAGGAGAATATTTGTAT	5023
ROR1-v3      ATAAATAAGCTGATTCAGAAGTTACATTTCTTAACTTTAGGTAGAGGAGAATATTTGTAT	4533


ROR1-v2      ------------------------------------------------------------	2305
ROR1-v1      CCTTTTGTTGCTATGCAACTGTTTAATGATGAAGCTTCAAACCACAATTTTGTATATCAT	5083
ROR1-v3      CCTTTTGTTGCTATGCAACTGTTTAATGATGAAGCTTCAAACCACAATTTTGTATATCAT	4593


ROR1-v2      ------------------------------------------------------------	2305
ROR1-v1      ATGACAATCAATTGTTTTCCAAGAATATTTATTATTTTAAGTACACAATTTCAGTGAATC	5143
ROR1-v3      ATGACAATCAATTGTTTTCCAAGAATATTTATTATTTTAAGTACACAATTTCAGTGAATC	4653


ROR1-v2      ------------------------------------------------------------	2305
ROR1-v1      TTGAGTTTTTCTAGGAGTCCCTTAAAGGGAAATTAGCATTCCATGAGCCAGTAAAATACC	5203
ROR1-v3      TTGAGTTTTTCTAGGAGTCCCTTAAAGGGAAATTAGCATTCCATGAGCCAGTAAAATACC	4713


ROR1-v2      ------------------------------------------------------------	2305
ROR1-v1      TACTCTGGTAAAACATTATTATGGTTAAAAAGTAAATTCAAGTTAGTTTTTTATAAAGAA	5263
ROR1-v3      TACTCTGGTAAAACATTATTATGGTTAAAAAGTAAATTCAAGTTAGTTTTTTATAAAGAA	4773


ROR1-v2      ------------------------------------------------------------	2305
ROR1-v1      CTATAACATTTATTTTAAACATTTTATAATAACTGAAAACATTAAAGTGAGCAAATGAAA	5323
ROR1-v3      CTATAACATTTATTTTAAACATTTTATAATAACTGAAAACATTAAAGTGAGCAAATGAAA	4833


ROR1-v2      ------------------------------------------------------------	2305
ROR1-v1      TTTCAAAACCACTTGTAATAATGTATTTTATAATCGCACTGTGATACTATATAACACAGT	5383
ROR1-v3      TTTCAAAACCACTTGTAATAATGTATTTTATAATCGCACTGTGATACTATATAACACAGT	4893


ROR1-v2      ------------------------------------------------------------	2305
ROR1-v1      CTCTTTTGTATTAAAATAGTATTTTTTTCACAAGCTGAAAAATATCTTTTCCTTTGTTGA	5443
ROR1-v3      CTCTTTTGTATTAAAATAGTATTTTTTTCACAAGCTGAAAAATATCTTTTCCTTTGTTGA	4953


ROR1-v2      ------------------------------------------------------------	2305
ROR1-v1      CAATGTTTTGTAAGATGACTTTATTTTCAGATCTTTTTCTCCTTTTTTGCACAAAACTAT	5503
ROR1-v3      CAATGTTTTGTAAGATGACTTTATTTTCAGATCTTTTTCTCCTTTTTTGCACAAAACTAT	5013


ROR1-v2      ------------------------------------------------------------	2305
ROR1-v1      GCTTATGGTTTGTGTCACAGAAGTGAAAATATATCTTTGCATTTTTATATCTGGTCTGTT	5563
ROR1-v3      GCTTATGGTTTGTGTCACAGAAGTGAAAATATATCTTTGCATTTTTATATCTGGTCTGTT	5073


ROR1-v2      ------------------------------------------------------------	2305
ROR1-v1      TTTCTTGTTTCCTTTGTTTTTAACTTGATATAGATTTTCTTCAAATATATAATGGCAATT	5623
ROR1-v3      TTTCTTGTTTCCTTTGTTTTTAACTTGATATAGATTTTCTTCAAATATATAATGGCAATT	5133


ROR1-v2      ------------------------------------------------------------	2305
ROR1-v1      TTCAGATATCTCACCTTACCATATCTTTCCTTATTTTCACTGCATGCATTTAATCACTGT	5683
ROR1-v3      TTCAGATATCTCACCTTACCATATCTTTCCTTATTTTCACTGCATGCATTTAATCACTGT	5193


ROR1-v2      ------------------------------------------------------------	2305
ROR1-v1      ATTACTTAATGTTTGATTTGTTATTATGGGCATTTCAAATAGGCAAGCATTGAATTGTAA	5743
ROR1-v3      ATTACTTAATGTTTGATTTGTTATTATGGGCATTTCAAATAGGCAAGCATTGAATTGTAA	5253


ROR1-v2      ------------------------------------------------------------	2305
ROR1-v1      TGACAAAAAGGCTATTTTATATTAAGGATATATGCATTTGTATTTCACACACCAGAGATG	5803
ROR1-v3      TGACAAAAAGGCTATTTTATATTAAGGATATATGCATTTGTATTTCACACACCAGAGATG	5313


ROR1-v2      -------------------------------------------------------	2305
ROR1-v1      ATATTAAACACTGATTATTTTATGCTGCTGTTTATTAAAAATGTTTACTATAAAA	5858
ROR1-v3      ATATTAAACACTGATTATTTTATGCTGCTGTTTATTAAAAATGTTTACT------	5362
```
